# *Ultrabithorax* modifies a regulatory network of genes essential for butterfly eyespot development in a wing sector-specific manner

**DOI:** 10.1101/2022.03.20.485072

**Authors:** Yuji Matsuoka, Antónia Monteiro

## Abstract

Nymphalid butterfly species often have a different number of eyespots in forewings and hindwings but how the hindwing identity gene, *Ultrabithorax (Ubx),* drives this asymmetry, is not fully understood. It is also unclear why eyespot serial homologs originated initially only in hindwings. To address these questions, we examined a three-gene regulatory network (GRN) for eyespot development in the hindwings of *Bicyclus anynana* butterflies, and compared it to the same network previously described for forewings. We also examined how *Ubx* interacted with each of these three eyespot-essential genes. We found similar genetic interactions between the three genes in fore and hindwings, but we discovered three regulatory differences: *Antennapedia* (*Antp*) merely enhances *spalt* (*sal*) expression in the eyespot foci in hindwings but is not essential for *sal* activation, as in forewings; *Ubx* up- regulates *Antp* in all hindwing eyespot foci but represses *Antp* outside these wing regions and*; Ubx* is regulating *sal* expression in a wing-sector specific manner, i.e., it does not affect *sal* expression in the wing sectors that contain eyespots in both fore and hindwings, but it positively regulates *sal* expression in the sectors that have hindwing-specific eyespots. These results suggest that *Ubx*, or its downstream targets, might have paid a role in the origin of eyespots, restricted initially to hindwings, via the positive regulation of *sal*, an essential gene for eyespot development. We propose that *Antp* got co-opted into the eyespot GRN at a later stage by coming under *Ubx* regulation. This Hox gene redundancy, together with a novel positive feedback loop between *sal* and *Antp*, might have allowed *Antp* to functionally replace *Ubx* in forewings and lead to the origin of forewing eyespots. Outside the eyespot focal regions, we discovered that *Ubx* is up-regulated by *Distal-less* along the wing margin. We propose a model for how the regulatory connections between these genes might have evolved to produce wing- and sector-specific variation in eyespot number.

## Introduction

Hox genes are general transcription factors that are often used to promote or repress the development of traits. When Hox genes are manipulated, traits might become reduced or disappear, or become enlarged. For example, the Hox gene *Ultrabithorax (Ubx)*, is responsible for repressing the growth of the dorsal appendages of the third thoracic segment of *Drosophila* and shaping them into the small haltere balancing organs (Lewis 1978). The removal of *Ubx* from halteres makes these appendages develop into large flight wings (Lewis 1978). Conversely, the large hindlegs of crickets (Mahfooz et al., 2007) also owe their appearance to *Ubx,* which functions as a growth promoting gene in this species; hindlegs become smaller when *Ubx* is down-regulated. So, Hox genes can function as either promotors or repressors of traits, depending on the trait and species in question.

Recently, however, we have described a system where Hox gene manipulations affect the same type of trait in different ways, depending on the location of that trait within the body. In *Bicyclus anynana* butterflies, CRISPR-Cas9 experiments targeting *Ubx* led to an expected homeotic transformation, i.e., hindwing wing patterns were modified into those of the forewing (Matsuoka and Monteiro, 2021). This homeotic transformation led both to the *enlargement* of some eyespots as well as the *disappearance* of other eyespots on the hindwing. Eyespots that were unique to hindwings, i.e., without a corresponding serial homolog in forewings, required *Ubx* to differentiate, and did not differentiate when *Ubx* was disrupted. Eyespots with a forewing serial homolog, on the other hand, were repressed in size by *Ubx*. These hindwing eyespots, which are naturally smaller than their counterparts on the forewing, enlarged to forewing size when *Ubx* was disrupted. This indicated that *Ubx* is both a size repressor as well as an essential gene for eyespot development, and these two functions vary with the location of the eyespot on the hindwing.

Similar experiments targeting the Hox gene *Antennapedia* also led to different effects on eyespots, depending on whether the eyespots were on the forewings or hindwings. Antp protein is present in the center of all eyespots, but *Antp* disruptions led to the disappearance of all forewing eyespots, but only to the disappearance of the white centers and to a size reduction of hindwing eyespots (Matsuoka and Monteiro 2021). This indicated that *Antp* is essential for eyespot development on forewings, but merely essential for the differentiation of the white central scales and for enlarging eyespots on hindwings.

In contrast to the effects of disruptions of these two Hox genes, CRISPR-Cas9 targeted disruptions of two other eyespot-associated genes, *Distal-less* and *spalt*, in different nymphalid species, showed each gene to be essential for the development of all eyespots, on both forewings and hindwings, regardless of the wing sector where the eyespot was found (Zhang and Reed, 2016; Connahs et al. 2019; Murugesan et al. 2022). This suggests that some genes have a global effect on eyespot development whereas the Hox genes have a more limited, wing or wing-sector specific role.

Recently, we studied how three of these eyespot-associated genes, *Antp, Dll* and *sal,* interacted with each other on the forewings of *Bicyclus anynana* butterflies (Murugesan et al. 2022). We discovered that *Dll* is upstream of both *sal* and *Antp*, and is required to up-regulate these genes in eyespot centers, during the larval stages. We also discovered that *sal* and *Antp* up-regulate each other. The output of this regulatory network, however, must be modified on the hindwing by *Ubx,* seeing how *Antp* no longer functions as an essential gene for eyespots to develop on this wing. In addition, *Ubx* is required to develop eyespots that are unique to hindwings, but it is unclear whether this gene interacts or activates any of the other three genes in those wing sectors.

Given that *Ubx* is essential for the development of some hindwing eyespots (Matsuoka and Monteiro 2021), we hypothesized that *Ubx* might interact with at least one of the other three essential genes, *Antp, Dll,* or *sal* in hindwing eyespots. Either functioning upstream or downstream of these genes. We also hypothesized that *Ubx* might have a distinct interaction with these genes in sectors where *Ubx* merely displays a repressive effect on eyespot size. To test these hypotheses, we first described patterns of gene expression for all four genes in hindwings and compared them to forewings. Then we generated mosaic crispants with the CRISPR-Cas9 system, targeting *Ubx*, *Antp*, *Dll* and *sal,* in turn. We followed the effects of these perturbations on the protein levels of the targeted gene, as well as the other three proteins on the hindwing, using double-immunostains. We focused our investigation on the larval stages of hindwing development when eyespot centers, also called the foci, are being differentiated. We examined interactions between the four genes specifically in the foci, as well as in other parts of the wing.

## Results

### Differences in expression pattern of Dll, Sal, Antp, and Ubx proteins between fore and hindwings

To test whether there were differences in the expression pattern of Dll, Sal, and Antp proteins between fore and hindwings, we examined larval wings using immunohistochemistry. The adult forewing has only two eyespots in M1 and Cu1 wing sectors whereas the hindwing has seven eyespots (Fig. 1). We found that the expression pattern of Dll and Sal proteins was quite similar in both fore and hindwings, even though wing shape and final eyespot number are different (Fig. 1). In early larval wings both Dll and Sal proteins showed a similar finger pattern of expression in both fore and hindwings, that ended at the center of nine potential eyespots on each wing (Fig. 1). In contrast, the expression pattern of Antp protein was different between fore and hindwings. Antp protein was expressed in seven eyespot foci in hindwings, never in the finger pattern, but was only expressed in four foci, from M1 to Cu1 eyespots, in forewings (Fig. 1). Later in larval development, Antp expression was retained in M1 and Cu1 foci (Figsup. 2, 3, and 4), but lost from the middle two wing sectors.

**Figure 1.**
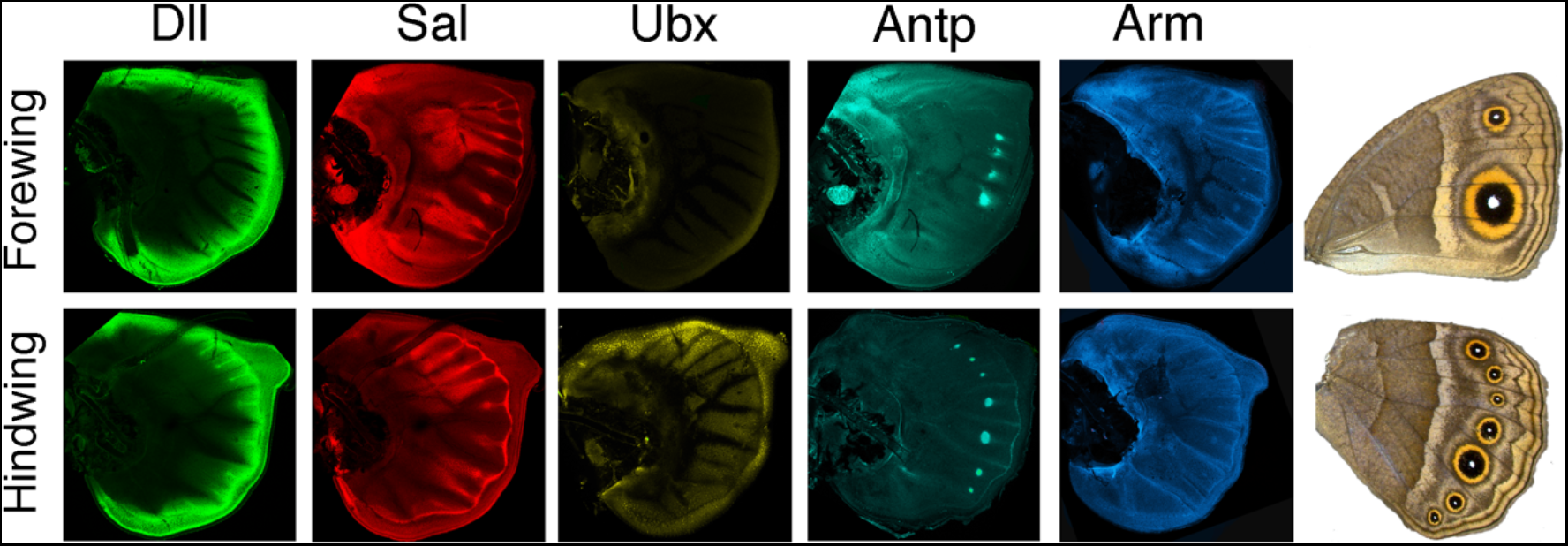
Differential expression pattern of eyespot associated genes between fore and hindwing. The Dll protein pattern is similar across both wings. Dll protein is present in the wing margin, along a finger pattern from the wing margin into nine wing sectors, and in the foci mapping to future eyespot centers. The Sal protein pattern is also similar in both wings. Sal shows a wing sector specific pattern of expression, with two outer domains, in the most anterior and most pesterir wing sectors, and two more central domains, one spanning wing sectors R2 to M3, and the other spanning A2 to A3 wing sectors (see Banerjee and Monteiro 2020). Fingers and foci pattern are similar to those of Dll. Ubx protein is ubiquitously present across the hindwing but was not detected in forewings. Ubx expression in the eyespot foci is stochastic. In late larval wings Ubx is sometimes up-regulated in the foci but in other wings not. Antp protein pattern is different between fore and hindwings. Antp is present only in four eyespot foci in the forewing, but in seven in the hindwing. Arm shows a similar pattern in eyespot foci as of Antp, but Arm is also expressed in the fingers leading to the foci.

To explore further differences between the eyespot GRN between fore and hindwings, we examined the expression pattern of Armadillo (Arm) protein, a mediator of Wnt signaling that is involved in eyespot development (Ozsu and Monteiro 2017). While Arm was expressed in all seven eyespot foci in hindwings, in forewings, Arm followed the dynamic Antp expression pattern, initially expressed in four foci and later only in two (Fig. 1, Figsup. 4). These stainings suggest that Dll and Sal proteins are not sufficient, on their own, to activate future eyespot centers on forewings; presence of Antp and Arm proteins, throughout the middle and late stages of larval wing development, might be necessary for the differentiation of an eyespot focus.

Ubx shows a typical protein expression pattern restricted to the hindwing, as previously shown (Fig. 1; Weatherbee et al. 1999, Tong et al. 2014). Ubx protein is ubiquitously present across the hindwing at the early larval stage. As the wing develops, Ubx protein disappears from the future eyespot centers in some wings, whereas in other wings, Ubx becomes over-expressed in these centers (Figsup. 5). These variations, which seem somewhat stochastic and not dependent on development time, affect all hindwing eyespots in the same way.

To examine the regulatory interactions between *Ubx*, *Dll, sal* and *Antp* in eyespot development in hindwings we used embryonic injections of Cas9 mRNA and single guide RNAs targeting one gene at a time.

### Genetic interaction between *Dll* and *sal*

We first examined the relationship between *Dll* and *sal* in hindwings (Fig. 2A). In *Dll* crispants with broad clones showing absence of Dll protein, Sal protein was broadly over-expressed in distal wing areas with no clear delineation of expression in the midline fingers and eyespot centers (Fig. 2B, Figsup. 6). This indicates that *Dll* is repressing *sal* in the distal part of the wing. In other *Dll* crispants, Dll null clones overlapping the eyespot foci and fingers disrupted Sal protein expression and resulted in no fingers and in a split eyespot focus (Fig. 2B-B’’’). In addition, Sal protein was not detected in the Dll-null clones in the chevron patterns along the wing margin (Figsup. 6), indicating that *Dll* is required for proper *sal* expression in chevrons along the wing margin, midline fingers, and eyespot centers.

**Figure 2.**
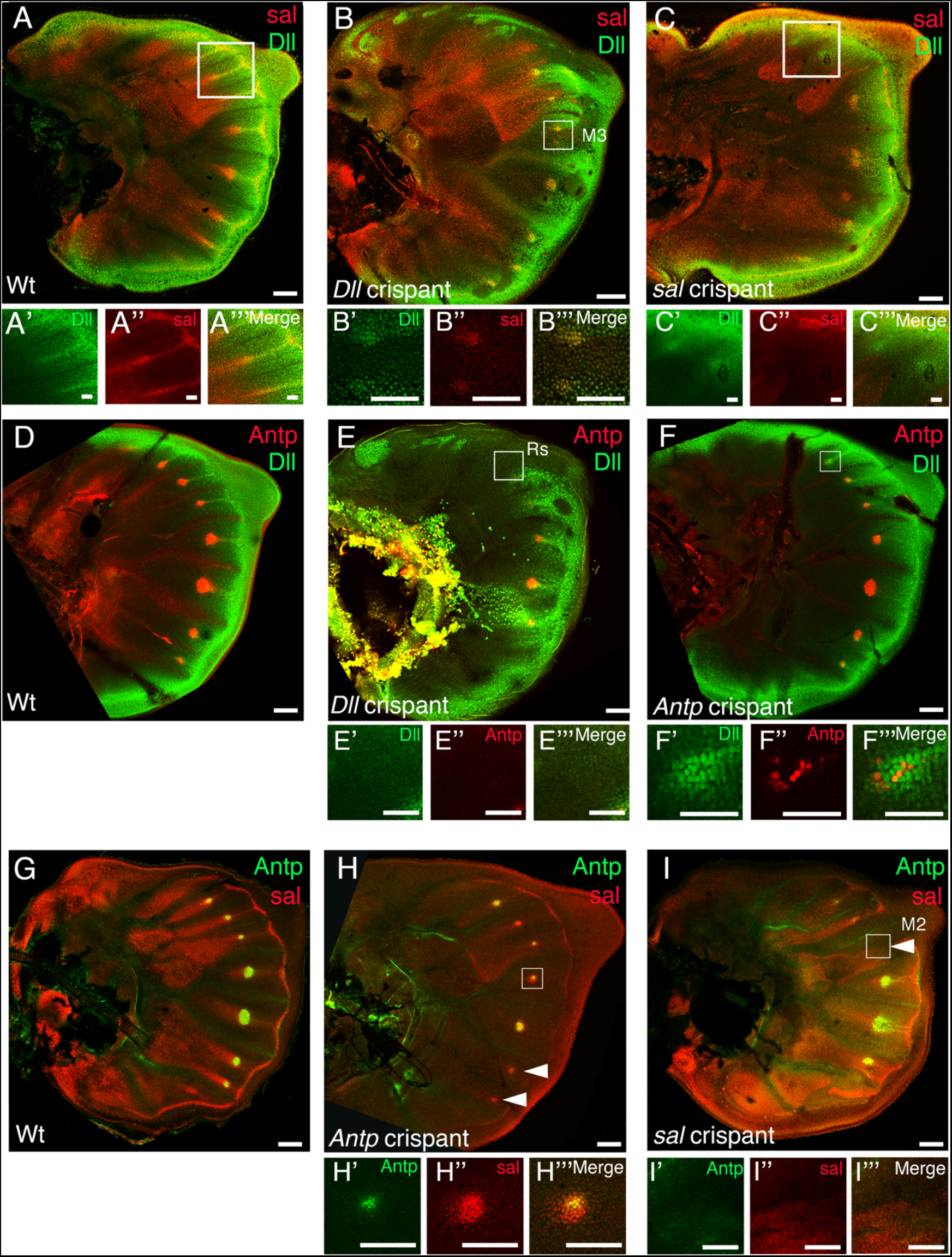
Regulatory interactions between Dll, sal, and Antp on the hindwing (A) Expression pattern of Dll and Sal proteins in a Wt hindwing. (B) Expression pattern of Dll and Sal in a *Dll* crispant hindwing. (C) Expression pattern of Dll and Sal in a *Dll* crispant hindwing. (D) Expression pattern of Dll and Antp in a Wt hindwing. (E) Expression pattern of Dll and Antp in a *Antp* crispant hindwing. (F) Expression pattern of Dll and Antp in a *Antp* crispant hindwing. (G) Expression pattern of Antp and Sal in a Wt hindwing. (H) Expression pattern of Antp and Sal in a *Antp* crispant hindwing. (I) Expression pattern of Antp and Sal in a *sal* crispant hindwing. Highly magnified regions are indicated with white squares. Arrowheads point to missing eyespot foci. Scale bar for whole wing images: 100 μm, for high magnification images: 50 μm.

We then examined whether *sal* also regulates *Dll*. *Sal* crispants with broad Sal-null clones showed normal Dll protein expression along the mid-line finger and wing margin (Fig. 2C-C’’’). However, eyespot centerswere split in two, when smaller Sal-null clones transected those cells (Figsup. 7). These results suggest that *Dll* is upstream of *sal,* as also observed in forewings (Murugesan et al. 2022), but *sal* is also regulating the focal expression of *Dll* in hindwings. Taken together, *Dll* appears to be an activator of *sal* in the eyespot centers, mid-line fingers, and chevrons along the wing margin, even though Sal protein starts to be visibly expressed in eyespot foci earlier than Dll protein (Figsup. 1). *sal* also regulates *Dll* in the foci. The regulatory relationship between these genes is distinct in the general distal wing region, where *Dll* is repressing *sal*.

### Genetic interaction between *Dll* and *Antp*

Next, we examined the relationship between *Dll* and *Antp* (Fig. 2D). In *Dll* crispants, Antp protein expression was lost in Dll-null clones (Fig. 2E-E’’’, Figsup.8), whereas in *Antp* crispants, Dll protein expression was not affected in Antp-null clones (Fig. 2F-F’’’, Figsup. 9). These results suggest that *Dll* up-regulates *Antp* in the hindwing eyespots as also observed in forewings (Murugesan et al. 2022). From the above results we identified *Dll* as an upstream regulator of both *Antp* and *sal* genes in hindwing eyespots, as also observed in forewing eyespots.

### Genetic interaction between *Antp* and *sal*

Next, we examined the relationship between downstream genes *Antp* and *sal* (Fig. 2G). In *Antp* crispants, Sal protein levels were dampened (but not lost) in Antp-null clones (Fig. 2H-H’’’, Figsup. 10), whereas in *sal* crispants, Antp protein levels were lost in Sal-null clones (Fig. 2I-I’’’, Figsup. 11). These results indicate that *sal* is essential to activate *Antp*, but *Antp* merely up-regulates *sal*, but is not essential for *sal* activation. The regulatory interaction between these two genes is different from the interaction observed in forewings, where *Antp* is an essential gene for *sal* activation (Murugesan et al. 2022). Taken together, the regulatory interactions involving two of the three genes are different between forewings and hindwings. To examine whether *Ubx* was involved in mediating these differences in regulatory interactions we extended our investigation to *Ubx*.

### Genetic interaction between *Ubx* and *Dll*

We first tested the specificity of the UbdA antibody, which is known to recognize both Ubx and AbdA proteins (Kelsh et al., 1994). In *Ubx* crispants, we found patches of hindwing cells that clearly lost all fluorescence (Fig. 3), indicating that the UbdA antibody detects Ubx proteins, and that Abd-A is not likely being co-expressed with Ubx on the hindwing.

**Figure 3.**
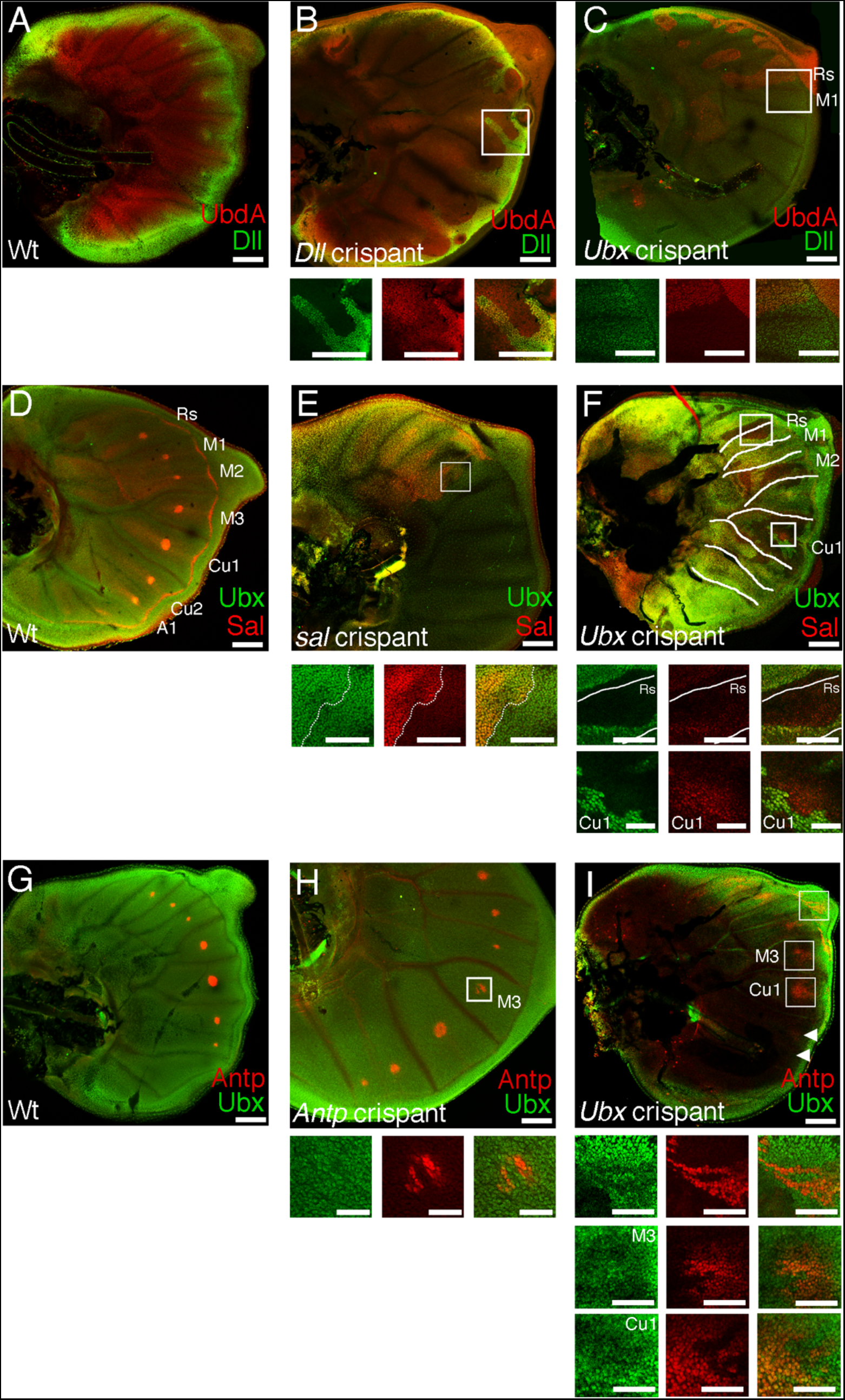
Regulatory interactions of Ubx with Dll, sal, and Antp on the hindwing (A) Expression pattern of Dll and UbdA protein in a Wt hindwing. (B) Expression pattern of Dll and UbdA in a *Dll* crispant hindwing. (C) Expression pattern of Dll and UbdA in a *Ubx* crispant hindwing. (D) Expression pattern of Ubx and Sal in a Wt hindwing. (E) Expression pattern of Ubx and Sal in a *sal* crispant hindwing. (F) Expression pattern of Ubx and Sal in a *Ubx* crispant hindwing. (G) Expression pattern of Antp and Ubx in a Wt hindwing. (H) Expression pattern of Antp and Ubx in a *Antp* crispant hindwing. (I) Expression pattern of Antp and Ubx in a *Ubx* crispant hindwing. Highly magnified regions are indicated with white squares. Scale bar for the whole wing images: 100 μm, for high magnification images: 50 μm.

We then examined the relationship between *Ubx* and *Dll* (Fig. 3A). In *Dll* crispants, Ubx protein expression was reduced in Dll null cells overlapping the wing margin (Fig. 3B, Figsup. 12). In *Ubx* crispants, the levels of Dll protein were not affected in both types of eyespot foci, those with forewing homologs and those present only on hindwings (Fig. 3C). Dll was also not affected in Ubx-null clones along the wing margin (Figsup. 13). However, it is unclear whether loss of *Ubx* activity affects the *Dll* pattern in the eyespot foci as Ubx expression is stochastic at the stage when we examined the relationship between the two genes. These results suggest that *Dll* is positively regulating *Ubx* along the wing margin, but *Ubx* does not regulate *Dll* expression.

### Genetic interaction between *Ubx* and *sal*

Next, we examined the relationship between *Ubx* and *sal* (Fig. 3D). In Sal-null cells, Ubx protein levels were not affected (Fig. 3E, Figsup. 14). However, in *Ubx* crispants, Sal protein levels were reduced, but not lost, in cells in the center of the wing (Figsup. 15). In the eyespot foci, the interaction of *Ubx* with *sal* was sector-specific and different between the sectors containing eyespots on both wings and sectors with hindwing-specific eyespots. Sal protein expression in the Rs and M2 eyespot foci was lost in Ubx-null clones (Fig. 3F, Figsup. 15D), whereas Sal protein expression in the Cu1 eyespot (an eyespot with a forewing serial homolog) was not affected in Ubx-null cells (Fig. 3F). These results suggest that *Ubx* is required to activate *sal* in the wing sectors with hindwing-specific eyespots, but not in the wing sectors with eyespots on both wings.

### Genetic interaction between *Ubx* and *Antp*

We next examined the relationship between *Ubx* and *Antp* (Fig. 3G). We found that Ubx protein levels were not affected in Antp null cells (Fig. 3H, Figsup. 16). However, *Ubx* regulates *Antp* in a distinct way in different wing regions. In the distal margin, surprisingly, Antp protein was up-regulated in clones of Ubx-null cells, with the level of ectopic Antp expression being comparable to the level observed in eyespot foci (Fig. 3I, Figsup. 17). In the foci, Antp protein expression was lost in both forewing serial homologs and hindwing-specific eyespot foci (Fig. 3I, Figsup. 17). These results suggest that *Ubx* is required for *Antp* activation in the eyespot foci, but it represses *Antp* along the wing margin.

In summary, we found that in hindwings, *Ubx* is required for *sal* activation in the hindwing-specific eyespot foci, and required for *Antp* expression in both kinds of eyespot foci. The main differences between the eyespot GRN in forewings and hindwings are that *Antp* is no longer essential to activate *sal* in hindwing foci, *Ubx* up-regulates *Antp* in all hindwing eyespots, and *Ubx* up-regulates *sal* in a sub-set of eyespots – those that are hindwing specific. Outside the foci, *Dll* is up-regulating *Ubx* expression along the wing margin, Ubx is up-regulating *sal* in more central wing regions, and repressing *Antp* along the wing margin and other areas of the wing.

## Discussion

### Differences in the differentiation of eyespot foci between fore and hindwings

*B. anynana* gradually differentiates two eyespot foci in larval forewings and seven foci in hindwings to produce a corresponding number of eyespots on the adult wings (Fig. 1). The genetic mechanism that limits the development of eyespots to the M1 and Cu1 wing sectors on the forewing is not understood. In this study we found that eyespot focal differentiation involves 1) Dll and Sal initially showing a similar protein expression pattern in both fore and hindwings across most wing sectors, 2) Antp and Arm proteins only becoming expressed in four of those sectors in forewings (M1, M2, M3 and Cu1), but in seven sectors in hindwings, and 3) the initially competent four middle forewing sectors becoming further reduced to two sectors (M1 and Cu1), with the disappearance of Antp and Arm proteins (as well as Sal, Dll, Engrailed and Notch proteins; Monteiro et al. 2013) from those sectors. A previous study identified a candidate locus, *Spotty*, that performs this last role – of repressing eyespot development from the middle M2 and M3 sectors in forewings – but its molecular identity is still unclear (Monteiro et al. 2013).

### Differences in the gene regulatory interaction of eyespot essential genes between fore and hindwings

The differences in the dynamic patterns of focal differentiation between forewings and hindwings are stemming from the exclusive presence of Ubx proteins in hindwings, as disruptions of this gene transform hindwing patterns into forewing patterns (Matsuoka and Monteiro 2021). These homeotic transformations include the removal of hindwing-specific eyespots, and the enlargement of two eyespots that have forewing homologs, indicating that Ubx interacts with these two types of eyespots in different ways. In this study, we examined how *Ubx* interacts with three other genes with an essential function in eyespot development in forewings (*Antp*) or both wings (*Dll* and *sal*), and how these interactions differ between forewings and hindwings and between hindwing sectors. We found six main results that are summarized in Fig. 4A-C: (1) *Dll* is required for both *Antp* and *sal* activation in hindwing eyespot foci, as also observed in forewings; (2) *sal* is required for *Antp* activation, as in forewings; (3) *sal* can impact *Dll* expression in the foci; (4) *Antp* up-regulates *sal* in eyespot foci but is no longer required for *sal* activation, as it is in forewings; (5) *Ubx* is required to activate *Antp* expression in all eyespot foci, and (6) *Ubx* is required for *sal* activation only in hindwing-specific eyespots. We will discuss these six results in turn.

**Figure 4.**
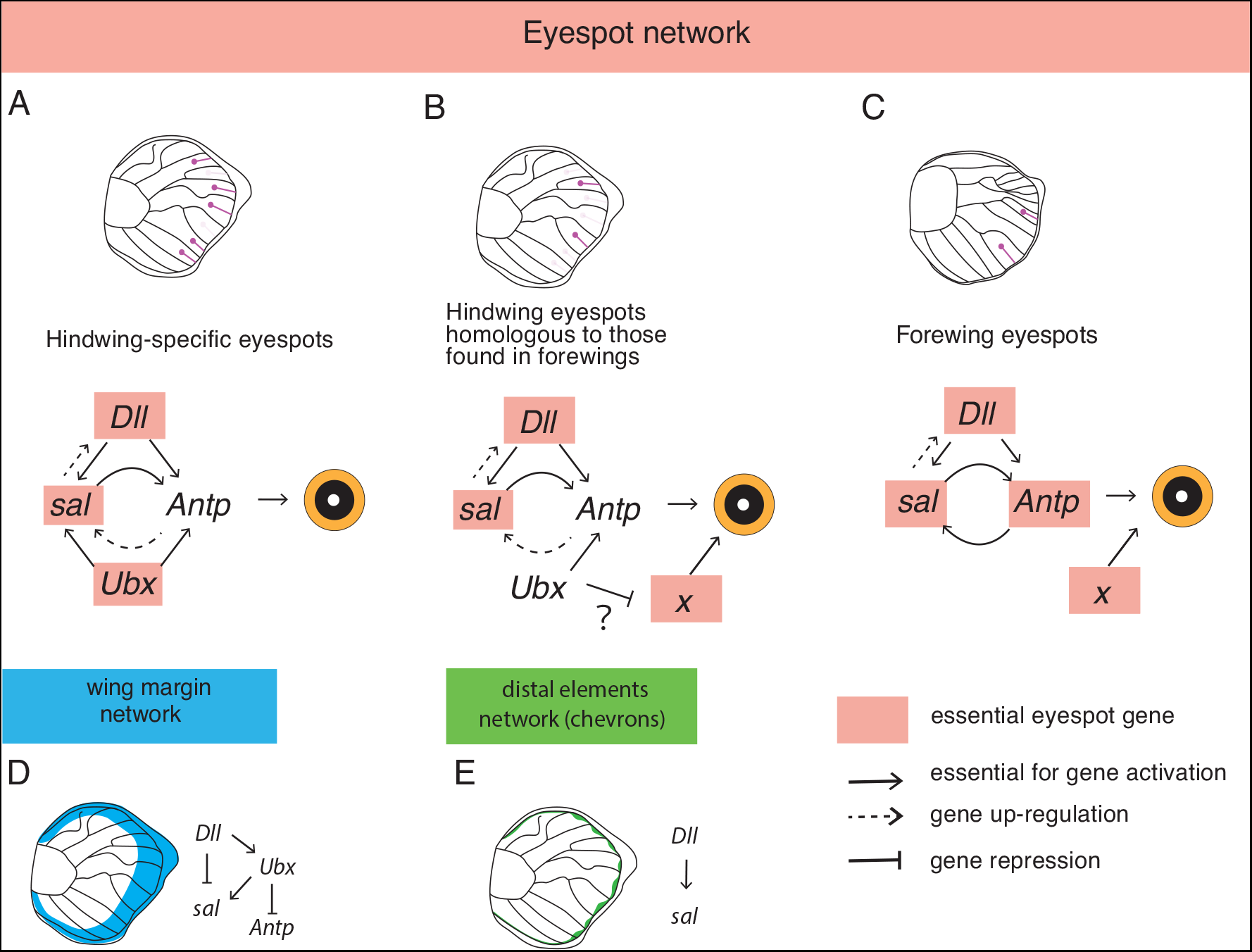
Schematic diagram of genetic interactions among Dll, sal, Antp, and Ubx on the hindwing (A) Genetic interactions in the focal region of the hindwing-specific eyespots. (B) Genetic interaction in the focal region of hindwing eyespots that have forewing homologs. (C) Genetic interaction in the eyespots on the forewing (from Murugesan et al. 2022). (D) Genetic interaction in the wing margin region. (E) Genetic interaction in the distal elements (chevrons) found in the wing margin.

Results (1) and (2) indicate that the eyespot GRN is largely similar between forewings and hindwings, that *Dll* is a top regulator, impacting both the expression of *sal* and *Antp*, and that *sal*, together with *Dll*, are essential for *Antp* activation.

Result (3) is novel as we observed that *sal* can also impact *Dll* expression in the foci, and split the foci into two, when specific Sal-null clones transect the foci, which was not observed in forewings (Murugensan et al. 2022). However, we speculate that this regulatory interaction might also be present in forewings because we observed split eyespot centers in both wings of adult *sal* crispants (Figsup. 18). In the previous study, the Sal-null clones examined covered the whole eyespot region, Dll expression in the fingers was not visibly affected, and the foci were not split (Murugesan et al., 2022). We propose that *sal* regulates *Dll* expression in the foci through the reaction-diffusion mechanism proposed for focus differentiation (Connahs et al., 2018). When this mechanism if disrupted, via loss of *sal*, the foci split apart. Further modeling of the reaction-diffusion mechanism, including *sal*, should be done in future.

Result (4) shows that while *sal* is still essential for *Antp* expression in hindwings, *Antp* merely up-regulates *sal* expression in the foci (Figsup. 10), rather than being essential for *sal* expression, as in forewings. We speculate that this regulatory difference between wings is, in part, due to redundant functions of *Antp* and *Ubx*, with *Ubx* also being involved in activating *sal* in some wing sectors, as discussed below.

Result (5), of *Ubx* being required to activate *Antp* expression in all eyespot foci, is novel. This positive regulation is contrary to what is normally observed in Hox genes, where more posterior Hox genes (such as *Ubx*) are often found to negatively regulate the more anterior Hox genes (such as *Antp*), at least in embryonic stages of insect development (Gummalla et al., 2014). While this negative cross-regulation is still observed overall on the hindwing, discussed below, it is not present in the focal cells. This indicates that the regulation of *Antp* by *Ubx* in the foci is novel.

In result (6) we found that *Ubx* regulates *sal* expression in a wing-sector specific manner. *Ubx* is essential for *sal* expression in the hindwing-specific eyespots but is not affecting *sal* expression in eyespots that have forewing homologs (Fig. 3F; Fig. 4). This result can help explain why *Ubx* disruptions led to loss of hindwing-specific eyespots alone, as *sal* is an essential gene for eyespot development, making *Ubx* also an essential gene for eyespot development in these wing sectors (Matsuoka and Monteiro 2021). How eyespots develop in the M1 and Cu1 hindwing sectors without *Ubx* input is unclear. In these sectors, *Ubx* is not up-regulating *sal* and is repressing eyespot size instead (Matsuoka and Monteiro 2021). We speculate that an unidentified gene (*gene X* in Fig. 4B, C), expressed in the M1 and Cu1 wing sectors, might be required to drive eyespot focus differentiation in these sectors, together with the other essential genes (*Dll* and *sal*).

*Antp* is no longer an essential gene in eyespot development in hindwings because disruptions of this gene merely result in loss of the white centers and in smaller eyespots (Matsuoka and Monteiro, 2021). This could be due to *Ubx* filling a partially redundant function with *Antp* in eyespot regulation, perhaps via the joint regulation of *sal.* This could happen due to the undifferentiated binding of both Hox genes to the same regulatory DNA sequences (Slattery et al., 2011). In forewings, *Antp* would be essential to activate *sal*, while in hindwings, *Ubx* could partially take over *Antp’s* role, making *Antp* no longer essential for eyespot foci differentiation. Finally, we discovered that *Ubx* and *Dll* are not regulating each other in the eyespot foci (Fig. 3B and 3C) These results suggest that at least two parallel inputs (*Ubx* and *Dll*) are necessary for the activation of *sal* and *Antp* in the hindwing-specific eyespots.

### Gene-regulatory interactions outside of the eyespot foci

In this study, we also examined regulatory interactions between the four genes outside the eyespot regions in hindwings of *B. anynana* (Fig. 4D, E). These data gave us extra insights about the GRN involved in general wing development and patterning, as this GRN is not well studied in insects other than *Drosophila*.

Here we show that *Dll* functions as both an activator as well as a repressor of *sal* outside of the foci (Fig. 2B, Figsup. 6, Fig. 4D, E). *Dll* functions as an activator of *sal* along the midline fingers and marginal chevrons (Fig. 4A, E). Previous computer simulations demonstrated that the finger and focal pattern of Dll is likely produced via a reaction diffusion mechanism that starts with Dll being uniformly expressed in the margin of the wing (Connahs et al. 2019). Our results suggest that eyespot foci, finger patterns, as well as margin chevrons might be the result of the same GRN, as also suggested recently in independent modelling work (Nijhout 2017). In the broader wing margin, however, outside the defined Sal stripe along the chevrons, *Dll* functions as a repressor of *sal* (Fig. 2B, and Figsup.6; Fig. 4D). To our knowledge, direct interaction of *Dll* and *sal* on the wing of insects has not been examined even in *Drosophila*. *Drosophila Dll* mutants show subtle changes in bristle formation along the wing margin that scarcely affect wing development (Campbell and Tomlinson, 1998). On the other hand, in a previous study, we found that loss of *Dll* affects wing shape, in addition to the loss of wing scales and eyespots in *B. anynana* (Connahs et al., 2018). Our data suggests that *Dll* is defining the distal limit of *sal* expression in butterfly wings, because *Dll* disruptions led to ectopic *sal* expression in the distal wing region, and resulted in deformed wings. These results suggest that the mechanism for wing margin development might be different between butterflies and flies.

Counteracting *Dll*’s repressive effects on *sal* in the wing margin, *Ubx* is positively regulating *sal* outside of the eyespot foci (Fig. 3F). A similar regulatory interaction is observed in *Tribolium*, where *Ubx* positively regulates *sal* in the flight wing (Tomoyasu et al., 2005). In the haltere of *Drosophila*, however, *Ubx* negatively regulates *sal* (Galant et al., 2002), so we speculate that this repressive function of *Ubx* is a derived feature of flies.

Interestingly, *Dll* also up-regulates *Ubx* in the distal wing margin, but *Ubx* does not regulate *Dll* (Fig. 3B and 3C; Fig. 4D). This finding is surprising as *Ubx* is usually understood as a modifier of the wing GRN, rather than being incorporated into the wing GRN and being itself modified by it. However, Ubx protein was not completely absent in *Dll* null cells (Fig. 3B), so *Dll* is not necessary for the activation of *Ubx* on the hindwing, but *Dll* increases *Ubx* expression along the margin. It is possible that this regulatory interaction between *Dll* and *Ubx* might have aided in the origin of eyespots initially restricted to hindwings, as discussed below.

It was surprising to discover that *Ubx* is repressing *Antp* in wing regions outside of the eyespot foci (Fig. 3I, Figsup.17). The same regulatory interaction is observed in *Drosophila* (Tsubota et al., 2008, Domsch et al., 2019). The up-regulation of *Antp* in *Ubx* crispants gives us an insight into the genetic mechanism behind the typical homeotic transformation in *Ubx* crispants, including that previously described for *B. anynana,* where hindwings were transformed into forewings (Matsuoka and Monteiro 2021). It has been believed that the forewing of most insects is a Hox-free state, and that *Ubx* gives hindwings their unique identity. However, in this study we found that Antp proteins were elevated in *Ubx* null cells (Fig. 3I, Figsup. 17), uncovering a repressive role of *Ubx* on *Antp* and the likely expression of *Antp* (albeit at low levels) in forewings as well (Fig. 4D). These results indicate that *Antp* is likely necessary for *B. anynana* forewing differentiation as has been recently shown for *Bombyx*, *Drosophila*, and *Tribolium* (Paul et al. 2021, Fang et al., 2022). In a previous study we also showed that Sal proteins were lost in *Antp*-null cells outside of the eyespot region in forewings (Murugesan et al. 2022), suggesting that Antp proteins are present in the wing at low levels, and are required for regulating *sal*. In addition, the shape of the forewing was slightly deformed in *Antp* crispants (Matsuoka and Monteiro, 2021). These results indicate that *Antp* is playing a role in forewing differentiation in *B. anynana*, and that butterfly forewings do not represent a Hox-free state.

Overall, we uncovered a partial gene regulatory network for wing development in *B. anynana* butterflies, and showed that even though the expression of wing patterning genes is highly conserved, their genetic relationship is slightly different from that of *Drosophila* and beetles.

### Possible genetic mechanism underlying eyespot origins and eyespot number evolution

Ancestral state reconstructions on a large phylogeny of ~400 genera suggested that eyespots first originated in four to five wing sectors on the ventral side of the hindwings of an ancestral lineage of nymphalid butterflies, before appearing on forewings and on dorsal sides of both wings (Oliver et al. 2014; Schachat et al., 2015). Recently, we proposed a possible function of *Antp* and *Ubx* as required genes for eyespot evolution (Matsuoka and Monteiro 2021). Here we provide further insight into the genetic mechanism that may have allowed the origin of eyespots on hindwings first, followed by eyespot origins on forewings.

Prior to the partial appendage GRN co-option event, proposed to have led to eyespot origins (Murugesan et al., 2022), nymphalid butterflies had no eyespots (Fig. 5A). We propose that after co-option of the appendage GRN, the genes *Dll* and *sal* gained a novel expression domain in the eyespot foci in fore and hindwings which, together with essential *Ubx* input restricted to hindwings, allowed eyespots to emerge in hindwings only (Fig. 5B). These eyespots might have originally lacked a white centre. The origin of forewing eyespots in satyrid butterflies might be connected to *Antp* having acquired a novel expression domain in eyespot foci, dependent on *Ubx*, which subsequently allowed *Antp* to take on the eyespot activating function of *Ubx* in forewings (Fig. 5C). In addition, the expression of *Antp* in eyespots might have aided the origin of the white centers (Fig. 5C). Eyespot number and size were further modified in each lineage and species probably through the introduction of mutations in genes such as *Spotty* (Fig. 5D). The Wt version of the *Spotty* gene represses eyespot development in the same two central wing sectors (M2 and M3) on both forewings and hindwings (Monteiro et al. 2007). *Ubx,* however, might be partially repressing *Spotty*, leading to the development of eyespots in those wing sectors in hindwings. *GeneX* might be involved in activating eyespots (perhaps independently) in M1 and Cu1 sectors in both wings, but on the hindwing the expression of this gene could be repressed by *Ubx,* leading to smaller M1 and Cu1 eyespots on the hindwing. Comparative work using functional gene expression studies is required to test these hypotheses.

**Figure 5.**
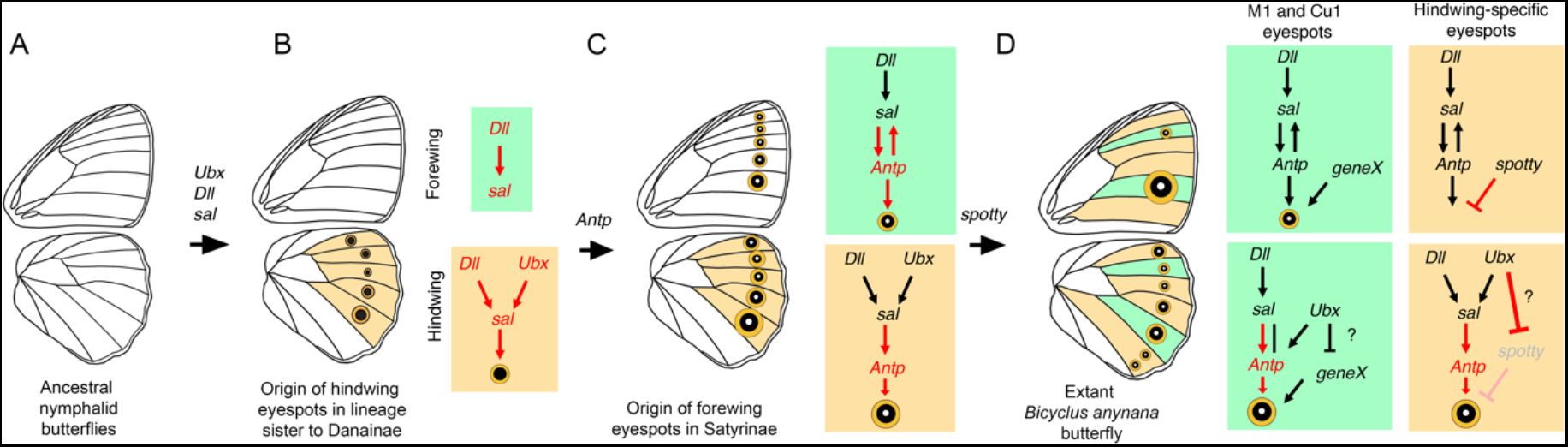
Proposed evolution of genetic interactions underling the origin of eyespots (A) Common ancestral nymphalid butterflies didn’t have eyespots on their wings. (B) The first eyespots likely appeared on the hindwing through the co-option of an appendage GRN. This might have led to the novel expression of *Dll* and *sal*, in the eyespot foci of hindwings alone, as Ubx is also essential for *sal* expression in most of the hindwing eyespots. (C) Once *Antp* was co-opted to the eyespot GRN, under *Ubx* regulation, the first eyespots would have orginated on the forewing, using *Antp* as the essential gene for *sal* activation in forewings. Eyespots might have also gained a distinct center at this stage. (D) The number and size of eyespot were likely modified in each lineage using sector-specific genes. The *spotty* locus led to the loss of eyespots in the M2 and M3 wing sectors on the forewing.

In conclusion, in this study we begun to address why a different number of eyespots develop on fore and hindwings of *B. anynana* butterflies by directly examining the regulatory interactions of a few eyespot essential genes, and their interaction with a hindwing-specific selector gene, *Ubx*. We uncovered part of a mechanism for wing sector-specific regulation of eyespot development directed by *Ubx*, and proposed a more detailed molecular mechanism to explain the hindwing-specific origin of eyespots. Future work, such as an analysis of the total RNA species present in each wing sector, as well as functional experiments across species, may further advance our understanding of the genetic mechanisms underlying eyespot number differences between wings and how eyespot evolution proceeded in nymphalid butterflies.

## Acknowledgements

This study was supported by the National Research Foundation (NRF), Singapore under its Investigatorship Programme (award NRF-NRFI05-2019-0006).

## Materials and method

### Butterfly husbandry

*B. anynana*, originally collected in Malawi, have been reared in the lab since 1988. Larvae were fed on young corn plans and adults on mashed banana. *B. anynana* were reared at 27°C and 60% humidity in a 12:12 light:dark cycle.

### sgRNA design

sgRNA target sequences were selected based on their GC content (around 60%), and number of mismatch sequences relative to other sequences in the genome (> 3 sites). In addition we picked target sequences that started with a guanidine for subsequent *in vitro* transcription by T7 RNA polymerase.

### sgRNA production

Template for *in vitro* transcription of sgRNA was made with PCR method described in Matsuoka and Monteiro, 2018. Forward primer contains T7 RNA polymerase binding site and sgRNA target site (GAAATTAATACGACTCACTATAGNN19GTTTTA GAGCTAGAAATAGC). Reverse primer contains the remainder of sgRNA sequence (AAAAGCACCGACTCGGTGCCACTTTTTCAAGTTGATAACGGACTAGCCTTATTTTAACT TGCTATTTCTAGCTCTAAAAC). PCR was performed with Q5 High-Fidelity DNA Polymerase (NEB) in 100 μl reaction volumes. After running PCR amplicon on a gel electrophoresis, PCR product was purified with Gene JET PCR purification kit (Thermo Fisher). In vitro transcription was performed with T7 RNA polymerase (NEB) using 500 ng of purified PCR product as a template for overnight. After removal of template DNA with DNase Ⅰ treatment, the RNA was purified with ethanol precipitation. The RNA was suspended to RNase-free water and stored at −80°C. sgRNA sequence were described in previous papers (Matsuoka and Monteiro, 2021, Murugesan et al., 2022).

### Cas9 mRNA production

Plasmid pT3TS-nCas9n (Addgene) was linearized with XbaⅠ and purified by Phenol/Chloroform and ethanol precipitation. *In vitro* transcription of mRNA was performed with mMESSAGEmMACHINE T3 kit (Ambion) using 1 μg of linearized plasmid as a template, and poly(A) tail was added to the synthesized mRNA by using Poly(A) Tailing Kit (Thermo Fisher). The RNA was purified with lithium-chloride precipitation, and then suspended to RNase-free water and stored at −80°C.

### Embryo microinjections

Butterflies were allowed to lay eggs on corn leaves for 30 min. We co-injected 0.5 μg/μl final concentration of sgRNA and 0.5 μg/μl final concentration of Cas9 mRNA into embryos within 2-3 h after egg laying. Eggs were sunk into PBS, and injection was performed inside the PBS. Food dye was added to the injection solution for visualization. Injected eggs were incubated at 27°C in PBS, and transferred on wet silk cotton on the next day, and further incubated at 27°C. After hatching, larvae were moved to corn leaves, and reared at 27°C with a 12:12h light:dark cycle and 60% relative humidity.

### Immunohistochemistry for embryos and wing tissues

Fifth instar and pupal wing tissues were dissected in PBS buffer under the microscope. The samples were fixed in 4% formaldehyde/Fix buffer (0.1 M PIPES pH 6.9, 1 mM EGTA pH 6.9, 1.0 % Triton x-100, 2 mM MgSO_4_) for 30 min on ice. The samples were washed with PBSTw for 3 times in every 10 min, and then the samples were kept in 5% BSA/PBSTw for 1 hour in block buffer, or further stored at 4°C.

The samples were replaced into a 5% BSA/PBSTw solution containing primary antibody, and incubated at 4°C overnight. We used a rabbit polyclonal anti-Dll (at 1:200, a gift from Grace Boekhoff-Falk), a mouse monoclonal anti-Antp 4C3 (at 1:200; Developmental Studies Hybridoma Bank), and a rabbit anti-*J. coenia* Ubx antibody (at 1:500; a gift from L. Shashidhara), and a rabbit anti-Sal (at 1:20,000; de Celis et al., 1999). For double staining, we added two primary antibodies in the same tube. The wings were washed 3 times with PBSTw, every 10 min. Then, the PBSTw was replaced with 5% BSA/PBSTw as a blacking reaction for 1 hour, and this solution was replaced with a 5% BSA/PBSTw solution with an appropriate secondary antibody (1:200), and incubated at 4°C for 2 hours. The wings were washed 3 times, every 10 min, and mounted in ProLong Gold mounting media. The images were taken under an Olympus FV3000 fluorescent microscope.

## Figure Legend

**Supplementary Figure 1. Expression pattern of Dll and Sal proteins during Wt wing development**

In early larval wings when trachea have not yet started to invade the lacunae in the wing pouch, Dll protein is present in the wing margin but not showing finger or foci patterns. Trachea invasion can be monitored as the dark features in the brightfield images. On the other hand, Sal already shows finger and foci patterns at the same stage. Sal also shows its wing sector specific expression. A similar pattern is observed in forewings and hindwings of same stage. When trachea start to invade the middle of the wing, Dll starts to show its expression in fingers and foci patterns. These patterns remain unchanged until the late larval stage, when Sal and Dll expression in the M2 and M3 forewing sectors disappear from the eyespot foci and fingers. Scale bar: 100 μm.

**Supplementary Figure 2. Expression pattern of Dll and Antp proteins during Wt wing development**

Antp protein expression is first detected at the stage when trachea spread to the middle of the forewing in the four wing sectors from M1 to Cu1. On the other hand, Antp expression gradually appears on the hindwing. First only in the M3 and Cu1 hindwing sectors, then in the M3 to A1 wing sectors. Expression of Antp in the rest of the wing sectors is not observed until the stage when trachea reach the wing margin. Scale bar: 100 μm.

**Supplementary Figure 3. Expression pattern of Antp and Sal proteins during Wt wing development**

The forewings of early larvae show Sal protein expression but not Antp. Antp expression appears in the wing at the stage when trachea starts to invade the lacunae. The pattern of both proteins is not changed until the stage when trachea almost reach the wing margin. After that stage, expression of these genes is retained only in the M1 and Cu1 foci. A similar temporal-spatial pattern of gene expression is observed on the hindwing.

**Supplementary Figure 4. Expression pattern of Antp and Arm proteins during Wt wing development**

Expression pattern of Antp and Arm proteins is synchronised for both wings. In the forewing, gene expression is initially detected in four wing foci, from M1 to Cu1, whereas later expression is retained only in the M1 and Cu1 foci. In the hindwing, the gene expression pattern for the seven eyespot foci gradually appears, as described above. Scale bar: 100 μm.

**Supplementary Figure 5. Expression pattern of Ubx and Antp proteins during Wt wing development**

In the forewing Ubx protein expression was not detected throughout wing development. In the hindwing, Ubx is ubiquitously expressed across the wing including in the eyespot foci. However, Ubx expression in the eyespot foci becomes stochastic from the stage when trachea reached the wing margin. In some wings expression is missing in the foci, and in other wings expression is at similar levels or at elevated levels relative to other wing cells. Scale bar: 100 μm.

**Supplementary Figure 6. Expression pattern of Dll and Sal proteins in *Dll* crispant hindwings**

(A, B) Wing margin region shows Dll-null cell clones and Sal becomes ectopically expressed in those cells. (C, D) Wing margin chevron (distal element) is mutated for Dll activity and Sal protein is lost in the Dll-null cells. (E) Wing margin region is mutated for Dll activity and Sal protein is ectopically expressed in the Dll-null cells. Clear contrast of Sal-positive and Sal-negative expression was observed. Highly magnified regions are indicated with white squares.

**Supplementary Figure 7. Expression pattern of Dll and Sal proteins in *sal* crispant hindwings**

(A) M3 eyespot focus is mutated for Sal activity, in which Dll expression is also affected. Wing margin distal element (chevron) is mutated for Sal activity, and Dll expression is not affected. (B) M3 eyespot foci is mutated for Sal activity, and Dll expression is affected in Sal-null cells. (C) Cu1 eyespot foci is mutated for Sal activity, and Dll (and Sal) expression is split into two foci. Highly magnified regions are indicated with white squares.

**Supplementary Figure 8. Expression pattern of Dll and Antp proteins in *Dll* crispant hindwings**

Whole wing is broadly mutated for Dll activity, and Antp protein expression is lost in multiple foci. Highly magnified regions are indicated with white squares.

**Supplementary Figure 9. Expression pattern of Dll and Antp proteins in *Antp* crispant hindwings**

Complete or partial absence of Antp activity in the foci does not impair Dll expression. Highly magnified regions are indicated with white squares.

**Supplementary Figure 10. Expression pattern of Antp and Sal proteins in *Antp* crispant hindwings**

(A) Cu1 eyespot is almost completely mutated for Antp activity. (B) M3 eyespot is almost completely mutated for Antp activity. (C) M3 eyespot is partially mutated for Antp activity. In all cases, Sal protein expression is only slightly diminished in Antp-null cells. Highly magnified regions are indicated with white squares.

**Supplementary Figure 11. Expression pattern of Antp and Sal proteins in a *sal* crispant hindwing**

Cu1 eyespot is partially mutated for Sal activity, and Antp protein expression is lost in Sal-null cells. Highly magnified region is indicated with white square.

**Supplementary Figure 12. Expression pattern of Dll and Ubda proteins in *Dll* crispant hindwings**

(A-C) The wing margin is broadly mutated for Dll activity, and UbdA expression is reduced in Dll-null cells. Highly magnified regions are indicated with white squares.

**Supplementary Figure 13. Expression pattern of Dll and Ubda proteins in a *Ubx* crispant hindwing**

Wing margin region is mutated for Ubx activity and Dll expression is not affected in Ubx-null cells. Highly magnified regions are indicated with white squares.

**Supplementary Figure 14. Expression pattern of UbdA and Sal proteins in *sal* crispant hindwings**

Several wing regions are mutated for Sal activity but Ubx expression is not affected in Sal-null cells. Highly magnified regions are indicated with white squares.

**Supplementary Figure 15. Expression pattern of Ubda and Sal proteins in *Ubx* crispant hindwings**

(A) Wing margin region is mutated for Ubx activity and Sal expression is slightly diminished in Ubx-null cells. (B) Central wing region is mutated for Ubx activity and Sal protein expression is slightly diminished in Ubx-null cells. (C) Wing cells of Sal expression domain just next to the Rs eyespot was mutated for Ubx activity. Sal expression likely not affected in the Ubx null cells. (D) Separate pictures of Fig. 3F. Sal expression in the M2 was lost in *Ubx* null cells.. Highly magnified regions are indicated with white squares.

**Supplementary Figure 16. Expression pattern of UbdA and Antp proteins in *Antp* crispant hindwings**

(A) Eyespot foci M2 to A1 lost all Antp activity, and Rs and M1 foci retain a bit of Antp activity. In all cases, Ubx expression is not affected in Antp-null cells. (B) M3 and A1 eyespot lost all Antp activity, and other eyespot partially lost Antp activity. Ubx expression is not affected in Antp-null cells in Cu1 eyespot foci. Highly magnified regions are indicated with white squares.

**Supplementary Figure 17. Expression pattern of UbdA and Antp proteins in *Ubx* crispant hindwings**

Distal wing regions are mutated for Ubx activity and Antp is upregulated in Ubx-null cells. Highly magnified regions are indicated with white squares.

## References

Banerjee T. D., Monteiro A. Molecular mechanisms underlying simplification of venation patterns in holometabolous insects. Development. 2020 147:dev196394.

Campbell, G., and Tomlinson, A. The roles of the homeobox genes *aristaless* and *Distal-less* in patterning the legs and wings of *Drosophila*. Development 1998 125:4483–93.

Chen P., Tong X. L., Li D. D., Fu M. Y., He S. Z., Hu H., Xiang Z. H., Lu C., Dai F. Y. *Antennapedia* is involved in the development of thoracic legs and segmentation in the silkworm, *Bombyx mori*. Heredity. 2013 111:182–8.

Connahs, H., Tlili, S., van Creij, J., Loo, T.Y.J., Banerjee, T., Saunders, T.E., and Monteiro, A. Disrupting different *Distal-less* exons leads to ectopic and missing eyespots accurately modeled by reaction-diffusion mechanisms. bioRxiv. 2017 https://doi.org/10.1101/183491.

Dion E., Monteiro A., Yew J. Y. Phenotypic plasticity in sex pheromone production in *Bicyclus anynana* butterflies. Sci Rep. 2016 6:39002.

Domsch, K., Carnesecchi, J., Disela, V., Friedrich, J., Trost, N., Ermakova, O., Polychronidou, M., and Lohmann, I. The Hox transcription factor Ubx stabilizes lineage commitment by suppressing cellular plasticity in *Drosophila*. Elife. 2019 8:e42675.

Fang, C., Xin, Y., Sun, T., Monteiro, A., Ye, Z., Dai, F., Lu, C. and Tong, X. The Hox gene *Antennapedia* is essential for wing development in insects. Development. 2022 149:dev199841.

Gummalla, M., Galetti, S., Maeda, R.K., and Karch, F. (2014). Hox gene regulation in the central nervous system of *Drosophila*. Front. Cell Neurosci. 23;8:96.

Kelsh, R., Weinzierl, R. O. J., White, R. A. H. and Akam, M. (1994). Homeotic gene expression in the locust *Schistocerca*: An antibody that detects conserved epitopes in ultrabithorax and abdominal‐A proteins. Dev Genet 15, 19–31.

Lewis EB. 1978. A gene complex controlling segmentation in *Drosophila*. Nature. 276:565–570.

Lewis D.L., DeCamillis M., Bennett R.L. Distinct roles of the homeotic genes *Ubx* and *abd-A* in beetle embryonic abdominal appendage development. Proc Natl Acad Sci U S A. 2000 97:4504–9.

Mahfooz N., Turchyn N., Mihajlovic M., Hrycaj S., Popadić A. *Ubx* regulates differential enlargement and diversification of insect hind legs. PLoS One. 2007 2:e866.

Matsuoka Y., Monteiro A. Melanin Pathway Genes Regulate Color and Morphology of Butterfly Wing Scales. Cell Rep. 2018 24:56–65.

Masumoto M., Yaginuma T., Niimi T. Functional analysis of *Ultrabithorax* in the silkworm, *Bombyx mori*, using RNAi. Dev Genes Evol. 2009 219:437–44.

Monteiro A., Prijs J., Bax M., Hakkaart T., Brakefield P. M. Mutants highlight the modular control of butterfly eyespot patterns. Evol Dev. 2003 5:180–7.

Monteiro A., Chen B., Scott L. C., Vedder L., Prijs H. J., Belicha-Villanueva A., Brakefield P. M. The combined effect of two mutations that alter serially homologous color pattern elements on the fore and hindwings of a butterfly. BMC Genet. 2007 11;8:22.

Monteiro A., Chen B., Ramos D. M., Oliver J. C., Tong X., Guo M., Wang W., Fazzino L., Kamal F. *Distal-less* regulates eyespot patterns and melanization in *Bicyclus* butterflies. J Exp Zool B Mol Dev Evol. 2013 320:321–31.

Monteiro A. Origin, development, and evolution of butterfly eyespots. Annu Rev Entomol. 2015 60:253–71.

Nagata T., Suzuki Y., Ueno K., Kokubo H., Xu X., Hui C., Hara W., Fukuta M. Developmental expression of the *Bombyx Antennapedia* homologue and homeotic changes in the *Nc* mutant. Genes Cells. 1996 1:555–68.

Nijhout, H.F. The Common Developmental Origin of Eyespots and Parafocal Elements and a New Model Mechanism for Color Pattern Formation. Diversity and Evolution of Butterfly Wing Patterns. pp 3–19.

Oliver J. C., Tong X. L., Gall L. F., Piel W. H., Monteiro A. A single origin for nymphalid butterfly eyespots followed by widespread loss of associated gene expression. PLoS Genet. 2012;8(8):e1002893.

Oliver J. C., Beaulieu J. M., Gall L. F., Piel W. H., Monteiro A. Nymphalid eyespot serial homologues originate as a few individualized modules. Proc Biol Sci. 2014 281. pii: 20133262.

Özsu, N., Monteiro, A. Wound healing, calcium signaling, and other novel pathways are associated with the formation of butterfly eyespots. BMC Genomics 2017 18, 788.

Saenko S. V., Marialva M. S., Beldade P. Involvement of the conserved Hox gene *Antennapedia* in the development and evolution of a novel trait. Evodevo. 2011 2:9.

Schachat S. R., Oliver J. C., Monteiro A. Nymphalid eyespots are co-opted to novel wing locations following a similar pattern in independent lineages. BMC Evol Biol. 2015;15:20.

Slattery M., Riley T., Liu P., Abe N., Gomez-Alcala P., Dror I., Zhou T., Rohs R., Honig B., Bussemaker H. J., Mann R. S. Cofactor binding evokes latent differences in DNA binding specificity between Hox proteins. Cell. 2011 147:1270–82.

Tong X., Hrycaj S., Podlaha O., Popadic A., Monteiro A. Over-expression of *Ultrabithorax* alters embryonic body plan and wing patterns in the butterfly *Bicyclus anynana*. Dev Biol. 2014 394:357–66.

Tong X. L., Fu M. Y., Chen P., Chen L., Xiang Z. H., Lu C., Dai F.Y. *Ultrabithorax* and *abdominal-A* specify the abdominal appendage in a dosage-dependent manner in silkworm, *Bombyx mori*. Heredity 2017 118:578–584.

Warren R. W., Nagy L., Selegue J., Gates J., Carroll S. Evolution of homeotic gene regulation and function in flies and butterflies. Nature. 1994 372:458–61.

Weatherbee S. D., Nijhout H. F., Grunert L. W., Halder G., Galant R., Selegue J., Carroll S. *Ultrabithorax* function in butterfly wings and the evolution of insect wing patterns. Curr Biol. 1999 9:109–15.

Xiang H., Li M. W., Guo J. H., Jiang J. H., Huang Y.P. Influence of RNAi knockdown for E-complex genes on the silkworm proleg development. Arch Insect Biochem Physiol. 2011 76:1–11.

